# Anatomical modeling of brain vasculature in two-photon microscopy by generalizable deep learning

**DOI:** 10.1101/2020.08.09.243394

**Authors:** Waleed Tahir, Sreekanth Kura, Jiabei Zhu, Xiaojun Cheng, Rafat Damseh, Fetsum Tadesse, Alex Seibel, Blaire S. Lee, Frédéric Lesage, Sava Sakadžié, David A. Boas, Lei Tian

## Abstract

**Objective and Impact Statement:** Segmentation of blood vessels from two-photon microscopy (2PM) angiograms of brains has important applications in hemodynamic analysis and disease diagnosis. Here we develop a generalizable deep learning technique for accurate 2PM vascular segmentation of sizable regions in mouse brains acquired from multiple 2PM setups. The technique is computationally efficient, thus ideal for large-scale neurovascular analysis.

**Introduction:** Vascular segmentation from 2PM angiograms is an important first step in hemodynamic modeling of brain vasculature. Existing segmentation methods based on deep learning either lack the ability to generalize to data from different imaging systems, or are computationally infeasible for large-scale angiograms. In this work, we overcome both these limitations by a method that is generalizable to various imaging systems, and is able to segment large-scale angiograms.

**Methods:** We employ a computationally efficient deep learning framework with a loss function that incorporates a balanced binary-cross-entropy loss and a total variation regularization on the network’s output. Its effectiveness is demonstrated on experimentally acquired in-vivo angiograms from mouse brains of dimensions up to 808×808×702 μm.

**Results:** To demonstrate the superior generalizability of our framework, we train on data from only one 2PM microscope, and demonstrate high-quality segmentation on data from a different microscope without any network tuning. Overall, our method demonstrates 10× faster computation in terms of voxels-segmented-per-second and 3× larger depth compared to the state-of-the-art.

**Conclusion:** Our work provides a generalizable and computationally efficient anatomical modeling framework for brain vasculature, which consists of deep learning based vascular segmentation followed by graphing. It paves the way for future modeling and analysis of hemodynamic response at much greater scales that were inaccessible before.

## 1 Introduction

The hemodynamic response to neural activation has become a vital tool in understanding brain function and pathologies [1]. In particular, measuring vascular dynamics has proved to be important for early diagnosis of critical cerebrovascular and neurological disorders, such as stroke and Alzheimer’s disease [2]. Existing tools for the measurement of cerebral vascular dynamics rely on functional imaging techniques, for example functional magnetic resonance imaging (fMRI), positive emission tomography (PET), and optical imaging [1, 3]. Importantly, mathematical models have been proposed for these neuroimaging methods, which provide valuable insight into the relation between the measured signals, and the underlying physiological parameters, such as cerebral blood flow, oxygen consumption, and rate of metabolism [4–7]. These mathematical models often require a topological representation of the blood vessels as a graph of spatially distributed nodes, connected via edges [5, 7]. These vascular graphs are usually estimated from two-photon microscopy (2PM) angiograms of the mouse brain [5], and segmentation of blood vessels is generally the first step in this process [8]. Vascular segmentation from cerebral 2PM angiograms, however, is a challenging task, especially for *in-vivo* imaging. Current state-of-the-art methods for this task [9, 10] suffer from limited computational speed, restricting their usefulness to only small-scale volumetric regions of the brain. Furthermore, due to rapid deterioration of measurement contrast with imaging depth in 2PM, these methods have been unable to demonstrate effective segmentation for vasculature deep beneath the brain surface. In this work, we address these limitations, and present a computationally efficient framework for 2PM vascular segmentation that allows us to effectively process much larger regions of the mouse brain compared to existing methods at significantly faster computation speed in terms of voxels segmented per second. Our method also demonstrates accurate segmentation for significantly deeper vasculature compared to the state-of-the-art.

Vascular segmentation involves assigning a binary label to each voxel of the input angiogram to indicate whether or not it is part of a blood vessel. This task is challenging, especially when dealing with 2PM angiograms, as the measurement contrast decreases sharply with imaging depth due to multiple scattering and background fluorescence [11]. Additional sources of measurement noise include motion artifact corruption during in-vivo imaging, large pial vessels on the cortical surface, and densely packed vasculature, making the segmentation task nontrivial. In the presence of these challenges, a number of techniques have been employed for vascular segmentation, including methods based on the Hessian matrix [12, 13], tracing [14], optimally oriented flux [15], and geometric flow [16]. However, in practice, these methods demonstrate limited segmentation quality [8].

In recent years, techniques based on deep learning have shown significant improvement over traditional methods for 2PM vascular segmentation [8–10, 17]. One of the first works in this line was done by Teikari et. al. [17], who presented a hybrid 2D/3D deep neural network (DNN) for the segmentation task. Their method utilized angiograms with shallow imaging depths (less than 100 *μm*), and were limited by computation speed. The segmentation quality was improved upon by Haft et. al. [10] by using an end-to-end 3D segmentation DNN. This model, however, similar to Teikari et. al., was also limited by slow computation, and required about one month to train on a dataset consisting of one annotated angiogram of dimensions 292 × 292 × 200 *μ*m. Damseh et. al. [18] improved upon this limitation, and were able to process much larger datasets with faster computation speed in terms of voxels segmented per second. Their framework used a DNN based on the DenseNet architecture [19], which processed the 3D angiograms by segmenting 2D slices one-by-one, and demonstrated better segmentation quality compared to previous methods. However, this DNN did not generalize with respect to various imaging setups, i.e. it performed good segmentation only for 2PM angiograms acquired on the same setup as the training data. Ideally, one would like to be able to segment angiograms from any 2PM microscope once the network has been trained. In order to overcome this limitation, Gur et. al. [9] recently proposed an unsupervised DNN based on the active contours method, and demonstrated improved generalization capability compared to supervised models [10, 17, 18], with faster segmentation speed. However, this method still suffers from excessive training and inference times, and high computational cost. Furthermore, lack of “supervised” information makes it difficult to segment deep vasculature, as severe noise corruption makes the task very challenging, even when using active contours [20]. These challenges limited its effectiveness to small-scale angiograms, with up to 200 μm imaging depth. Therefore, there is a need for a vascular segmentation method, that is not only able to generalize to different 2PM imaging setups, but is also fast and computationally efficient, to cope with the processing needs of large-scale angiograms.

In this work, we propose a novel deep learning method for vascular segmentation of cerebral 2PM angiograms, that overcomes the afrementioned limitations of existing techniques, and demonstrates state-of-the-art segmentation performance. Our contribution here is three-fold. First, we present a novel application of a total variation (TV) regularized loss function for 2PM vascular segmentation. The proposed loss function combines the “supervised” information from training data acquired on a single imaging setup, with an “unsupervised” regularization term that penalizes the total variation of the DNN output [21]. This regularization encourages piece-wise continuity in the final segmentation, and improves generalization ability of the trained DNN to different imaging setups, without the need of excessive training data, transfer learning [22], or domain transfer [23]. The TV-regularized loss also makes the DNN significantly more robust to mislabeled ground-truth annotations. This is particularly useful for large-scale 2PM vascular angiograms where significant noise in deep vasculature makes precise ground-truth annotation very challenging, even for human annotators, making ground-truth data prone to mislabeling. The TV penalty also imparts inherent denoising capability to the trained network, eliminating the need for any post-processing. Our second contribution is a novel pre-processing method, which aids in generalization by making the histogram of an arbitrary test angiogram similar to that of training data, in addition to reducing its noise. We demonstrate the effectiveness of this preprocessing method to improve segmentation quality, not only with our proposed method, but also with some existing 2PM vascular segmentation techniques. Our third contribution is the novel application of an extremely lightweight, end-to-end 3D, DNN for 2PM vascular segmentation, which is able to demonstrate an order of magnitude faster segmentation, compared to the current state-of-the-art [9], in terms of voxels segmented per second. This enables us to perform segmentation on significantly larger regions in several mouse brains, thus enabling large-scale in-vivo neurovascular analysis. To illustrate this unique capability, we demonstrate segmentation on a 808× 808× 702 μm volume in less than 2.5 seconds.

Following vascular segmentation from our DNN model, we perform graph extraction on the binary segmentation map using a recently developed method based on the Laplacian flow dynamics [18]. Importantly, we show that our segmentation results in better graph modeling of the vasculature across large volumes compared to other segmentation techniques.

Overall, we present a new high-speed and computationally efficient anatomical modeling framework for the brain vasculature, which consists of deep learning based vascular segmentation followed by graphing. Our work paves the way for future modeling and analysis of hemodynamic response at much greater scales that were inaccessible before. To facilitate further advancements in this field of research, we have made our DNN architecture, dataset, and trained models publicly available at https://github.com/bu-cisl/2PM_Vascular_Segmentation_DNN.

## 2 Results

### System Framework

The deep learning based vasculature anatomical modeling pipeline is shown in Figure 1(A). This modular framework takes 2PM angiograms of live mouse brain as the input, performs segmentation of blood vessels using a novel 3D DNN, and finally extracts a vascular graph from the network’s prediction. The DNN [Fig. 1(B), S7], detailed in section 4.3, is of critical importance in this pipeline and is our primary contribution, along with the novel application of a TV-regularized loss function for 2PM vascular segmentation, detailed in section 4.4. This network is first trained to minimize the discrepancy between manually annotated ground truth segmentation, and it’s own prediction [Fig.1(B)]. During this training process, the network is exposed to challenging regions in 2PM angiograms in order to improve its vessel recovery from poor quality images. Some examples of such regions include deep 2PM measurements with low signal contrast [Fig.2(A)], and pial vessel occlusions[Fig.2(A), red circle]. In addition, we use large input angiogram patches of size 128 × 128 × 128 voxels, in conjunction with a network optimized for computation speed, allowing us faster segmentation on significantly larger angiograms compared to state-of-the-art methods [9, 10]. Once trained, this network provides segmentation of 2PM angiograms in a feed-forward manner [Fig.1(C)] that outperforms the state-of-the-art methods as detailed below.

**Figure 1:**
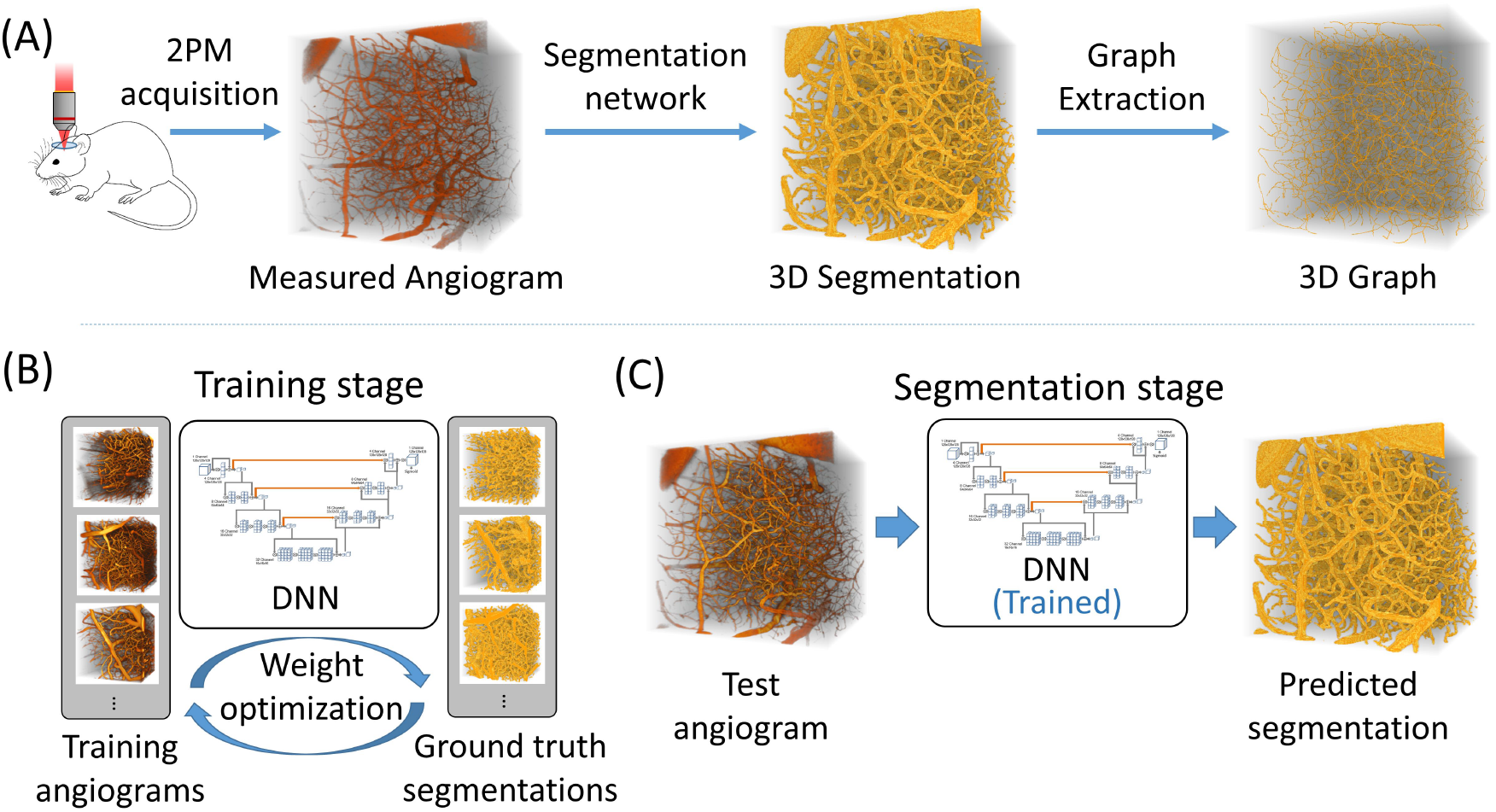
Framework for vascular modeling. (A) Two-photon microscopy (2PM) is used to acquire cerebral angiographic data on a live specimen via in-vivo imaging. This is followed by binary vascular segmentation of the 2PM angiogram. Finally, the 3D graph of the vasculature is computed from the segmentation map. In this paper, we present the segmentation method in detail, which is able to process large-scale 2PM angiograms. (B) A deep neural network (DNN) is used for segmentation which is first trained using annotated angiograms. During this process, the network weights are iteratively adjusted for accurate vessel segmentation. (C) After training is complete, the optimized network can be used in a feed-forward manner for segmentation on unseen angiograms.

**Figure 2.**
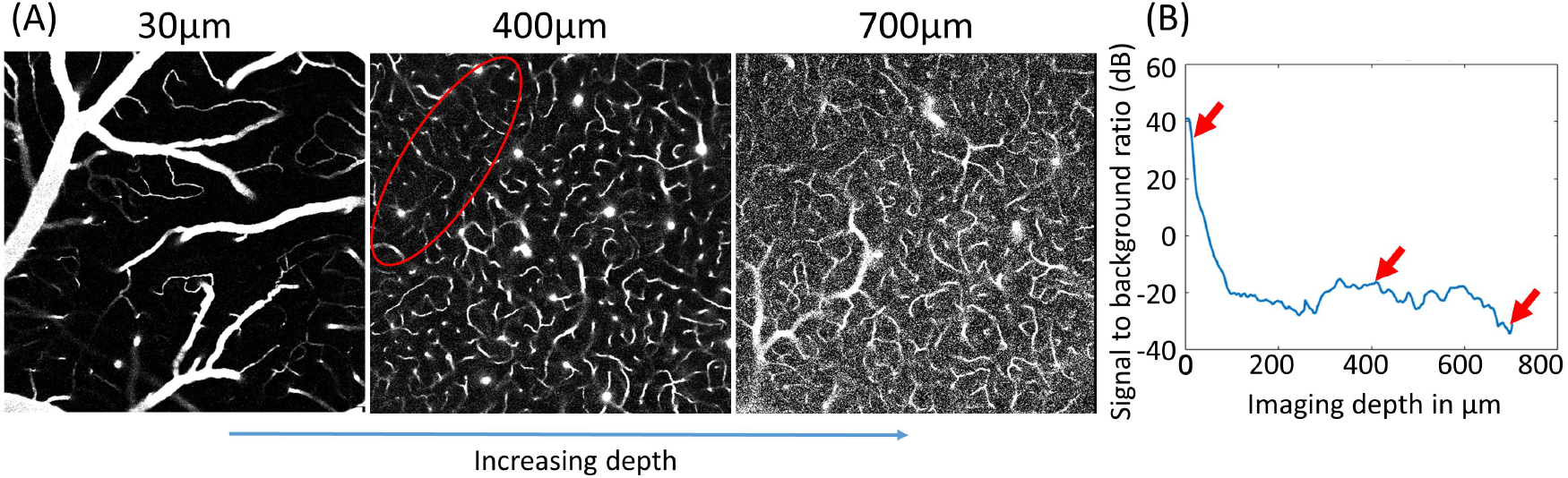
Deterioration of two-photon microscopy signal with imaging depth. (A) Visual contrast decreases for deeper vasculature due to loss of illumination focus with increased imaging depth, and higher background fluorescence due to increased laser power. Large pial vessels on the surface cast shadows underneath, as shown by the encircled region, making vessel detection challenging. (B) The signal to background ratio (SBR) of the angiogram decreases rapidly going deeper into the brain tissue.

### Segmentation Performance Analysis

To evaluate our segmentation approach, we first visually compare the predicted vessel segmentation from our DNN with the ground truth, a traditional Hessian matrix approach [13], and a recently developed DNN model [8] in Figure 3. Our method outperforms both these techniques in terms of segmentation quality, especially for vessels deep beneath the cortical surface. The Hessian matrix approach identifies tubular structures by an enhancement function based on Hessian eigenvalues. While it recovers most of the vessels closer to the surface, it preforms poorly in this regard for deeper vessels, due to significantly higher measurement noise in the angiogram, which makes it difficult to distinguish between the vessels, and the surrounding noisy background. Rafat et. al. [8] use a DenseNet architecture [19] to perform vascular segmentation in a 2D slice-wise manner. Although their network performs better than the Hessian approach, it also suffers from bad segmentation for deeper vessels, and ignores 3D context due to the slice-wise processing. A significant advantage that the proposed method possesses, compared to these techniques, is the TV-regularized loss function, which penalizes the variation of the DNN output, thus imparting inherent denoising capability to the network, and improving its segmentation for deep vessels. In addition, the proposed DNN also performs end-to-end 3D processing of data, which takes into account the 3D context of vasculature. Thus, the segmentation from our method maintains greater overlap of the prediction and ground truth compared to other methods, out to 606*μ*m [Fig. 3(B)-(D)]. These results indicate a 3× depth improvement using our approach over the current state-of-the-art methods. Since these large imaging depths also exhibit poor signal-to-background ratios (SBR) [Fig. 2], our DNN model also provides visually superior performance under low signal contrast imaging conditions. As discussed below, we quantify these improvements using a comprehensive set of metrics to holistically evaluate the vascular segmentation performance.

**Figure 3.**
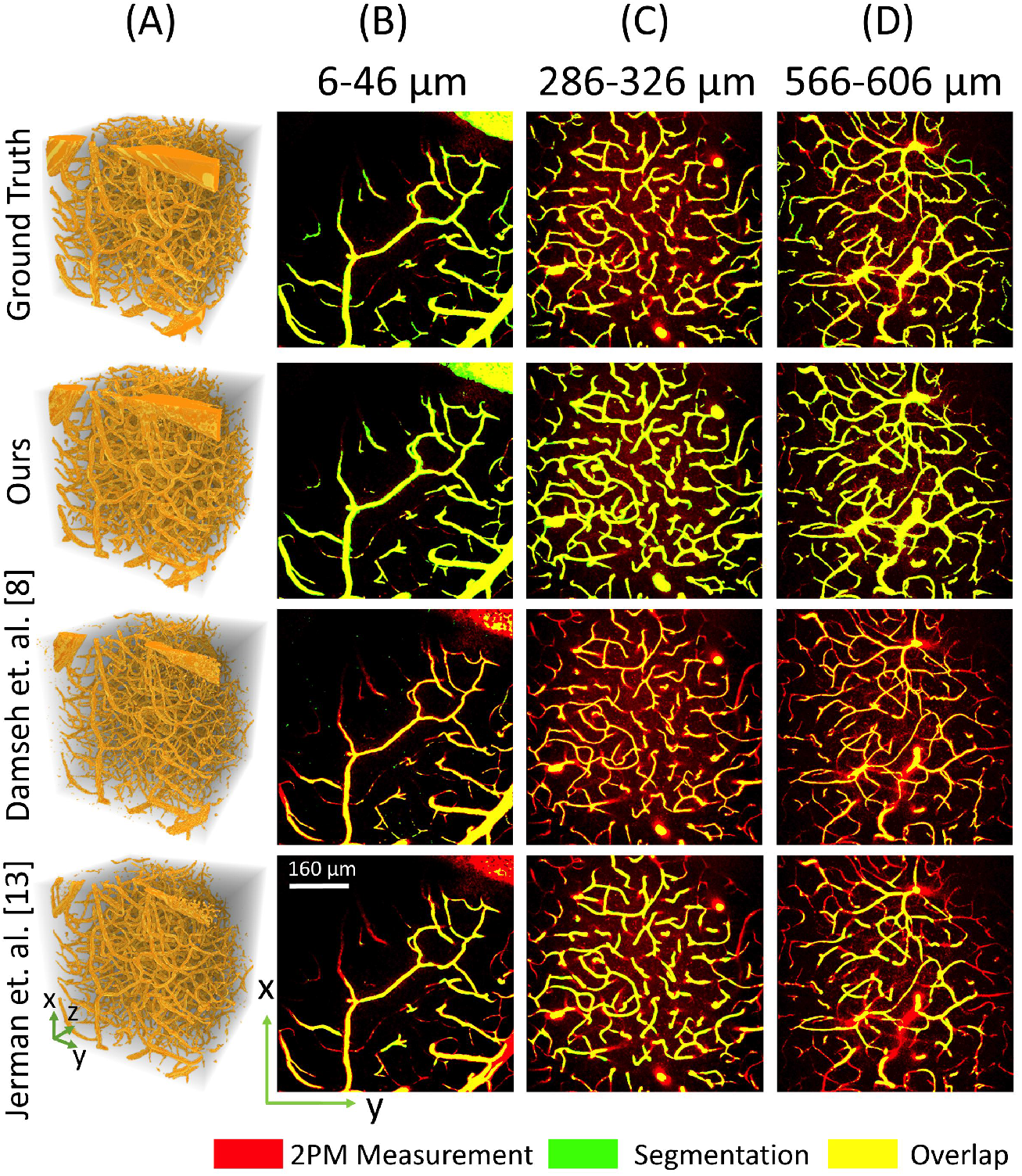
Large-scale 2PM vascular segmentation. (A) 3D renderings of segmentation on the test angiogram. (B-D) Maximum intensity projections (MIPs) of binary segmentation overlaid on 2PM measurement. Each MIP represents 40 *μ*m physical depth, and 20 discrete slices along the z-axis. MIPs for three different depths ranges are presented to show the effect of axial depth on segmentation performance. We demonstrate good segmentation for vasculature up to 606μm, despite significant increase in background noise associated with 2PM.

Overlap-based metrics are the most widely used metrics to evaluate vessel segmentation algorithms, which are computed based on analyzing the voxel overlap between the ground truth and the prediction. For example, sensitivity and specificity represent the respective percentage of the foreground and background voxels that are correctly recovered in the prediction. The Jaccard index computes the intersection over the union of the prediction and the ground truth, representing similarity based on percentage overlap. The Dice index is very similar to the Jaccard index and the two are strongly positively correlated. Generally, such metrics only compare the physical overlap between the ground truth and the predicted segmentation without considering the underlying morphological shapes of the object [24]. This factor makes overlap-based metrics ill-suited for delimiting complex boundaries like blood vessels, since they will preferentially correct larger vessels occupying more of the volume while ignoring the smaller, yet important capillaries in the vasculature. In addition, these metrics suffer from inherent biases towards one segmentation class or the other. For example, the Jaccard index, the Dice coefficient, and sensitivity are insensitive to true negative predictions, making them primarily indicative of positive class performance. On the other hand, specificity is insensitive to true positive predictions, thus primarily indicative of negative class performance. Accuracy is subject to class imbalance, i.e. when one type of class labels are significantly more abundant than the rest, accuracy becomes more indicative of the abundant class. This problem is particularly prevalent in vascular segmentation [25]. As an example, our manually segmented 2PM angiograms contain more than96% background tissue voxels and less than 4% foreground vessel voxels, indicating an imbalance ratio of more than 24.

To overcome these shortcomings, we further quantify our DNN performance using the correlation-based Matthew’s correlation coefficient (MCC) metric, two morphological similarity based metrics, namely Hausdorff distance (HD), and Modified Hausdorff distance (MHD), and a graph-based metric length correlation (LC). MCC is particularly suited for highly imbalanced data [26] since it is unbiased towards any class and gives the same value between −1 and 1, even when negative and positive classes are swapped. A score of 1 means perfect correlation, 0 means uncorrelated and the classifier is akin to random guessing, −1 means perfect negative correlation. HD measures the extent of morphological similarity between the prediction and ground truth, i.e. how visually similar their shapes are. This metric is suitable for data involving complex contours e.g. blood vessels [24]. MHD is a variant of HD, and is more robust to outliers and noise. LC is a graph-based metric, which we derive from the length metric in [27], and is specifically suitable for vascular segmentation. It measures the degree of coincidence between the predicted and the ground truth segmentations in terms of the total length. Since accurate graph extraction is the eventual goal for our segmentation pipeline, LC is a particularly well suited metric for comparison.

Quantitative evaluation of our DNN segmentation on (unseen) testing data is presented in Fig.4(A,B). Our method demonstrates the best overall segmentation, especially with respect to non-overlap-based metrics. In addition to providing both qualitatively and quantitatively improved vessel segmentation, our method also provides ≈ 10× faster voxel-per-second segmentation than the current state-of-the-art [9] as shown in Fig.4(C). Faster computation speed played an important role in enabling our DNN to train on our large-scale dataset within a reasonable time, and has potential applications for real-time segmentation.

**Figure 4.**
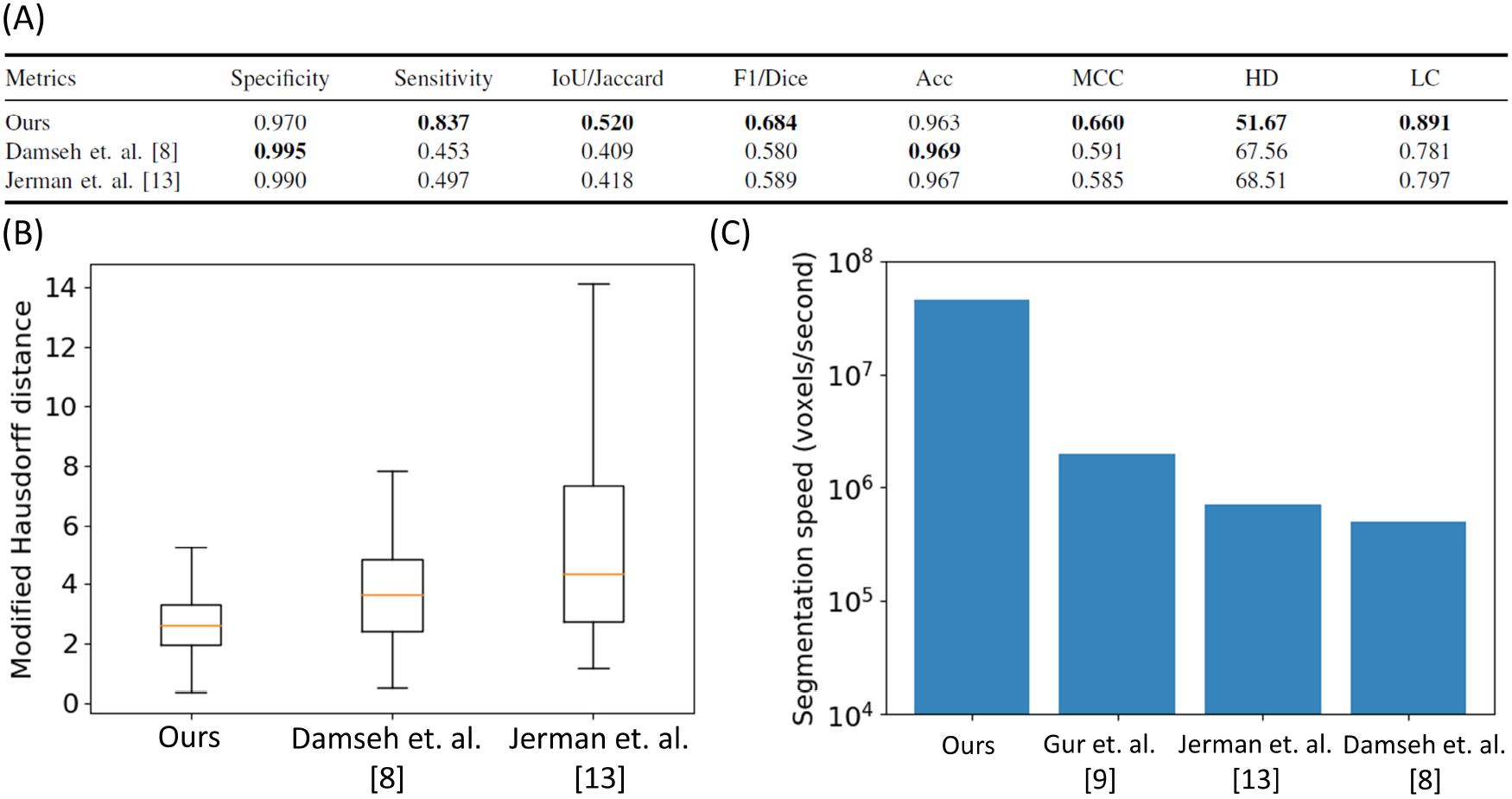
Quantitative evaluation of segmentation performance. (A) While we do present traditional metrics for segmentation comparison, e.g. sensitivity, specificity, Jaccard index, F1 score and accuracy; we also include other metrics arguably better suited for vascular segmentation including Matthew’s correlation coefficient (MCC), Hausdorff distance (HD), and length correlation (LC). Our method provides best overall performance on the test dataset, supporting the qualitative results in Fig.3. (B) Here we compare the slice-wise modified Hausdorff distance (MHD) between the methods of interest. Our method outperforms other techniques with considerably smaller mean and standard deviation of the slice-wise MHD. (C) In terms of number of voxels segmented per second, our method is about 10x faster than the state-of-the-art [9], making it suitable for large scale and real-time applications.

### Generalization to a new 2PM imaging system

Existing segmentation methods based on supervised learning [8, 10, 17] have not demonstrated the ability to generalize across various 2PM imaging setups, and the same setup is used to acquire both the training and testing data. In some cases, even the same 3D angiogram is divided into both training and testing sets [10]. Training a DNN with the ability to generalize over different setups is challenging due to inter-microscope variability. However, it is highly desirable to have a segmentation method which is independent of the acquisition hardware. One possible solution is to train a supervised DNN with annotated data from various imaging setups, because having such diversity in the training set is known to improve generalization performance. This however is difficult to achieve, since manually annotating ground truth for many large-scale angiograms from various setups is impractical due to the prohibitive labor cost. This is also why very few such large-scale annotated datasets are publicly available. Limited data availability makes it difficult to effectively train a purely supervised learning model. Supervised methods are also susceptible to possible mis-annotations, e.g. due to human error, and this problem is particularly amplified for large-scale datasets [21]. Another possible approach for achieving generalization is to train an unsupervised DNN, as by Gur et. al [9]. However, unsupervised methods discard annotated data altogether, and this may be detrimental in challenging regions of low contrast like deep vasculature, and areas under large pial vessels, where ground-truth annotations might be able to guide the training process. Here we present a DNN with a TV-regularized loss function, which combines the benefits of supervised learning with an unsupervised regularization term that penalizes the total variation of the DNN output, to demonstrate state-of-the-art generalization performance. Such a regularized learning scheme is not as susceptible to mis-annotations as supervised methods, and is demonstrated empirically in Fig. S1. In addition, unlike unsupervised methods, our network is able to incorporate expert annotated data for training, which is especially beneficial for low contrast and high noise regions in deep vasculature. After being trained on data from only one imaging setup, we demonstrate that our network is able to demonstrate good segmentation on data from another 2PM microscope without any retraining or network fine tuning [Fig.5].

**Figure 5.**
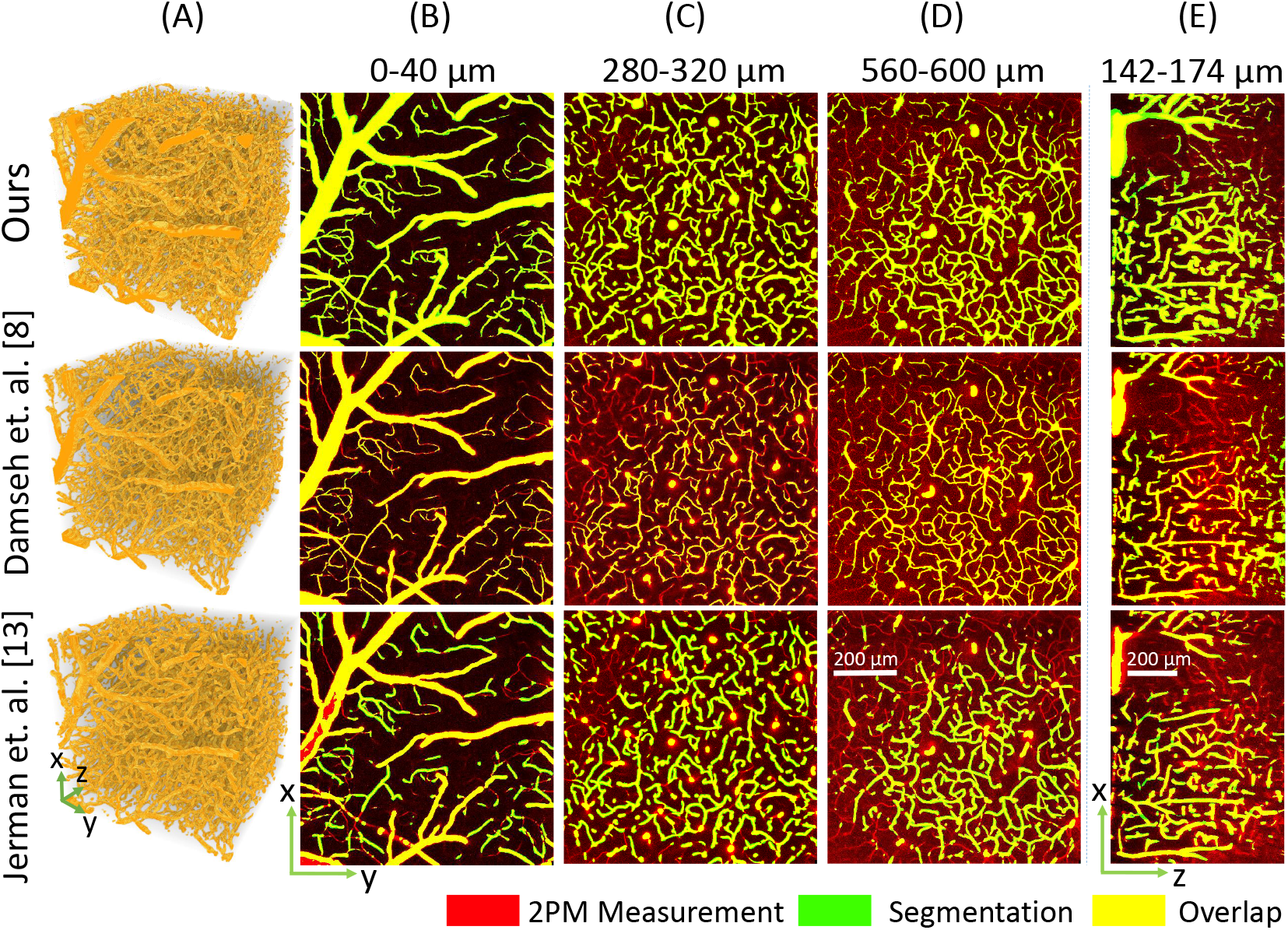
Generalization capability of our segmentation method. Our DNN is optimized for robustness to inter-microscope variability. We train our DNN on data from one 2PM setup and test on an angiogram acquired on a different setup, demonstrating good segmentation quality. (A) 3D renderings of the segmentation maps. (B-D) Maximum intensity projections (MIPs) of vascular segmentation overlaid on 2PM measurement, shown for lateral x-y cross sections. Each MIP represents 20 discrete slices along the z-axis. Our method has good qualitative performance with well connected vasculature and apt segmentation for both large and small vessels, demonstrating its ability to generalize to other 2PM imaging setups without retraining. For comparison, the supervised learning method by Damseh et al. [8] is unable to generalize well, and the resulting segmentation is not well connected. (E) MIPs for longitudinal x-z cross sections, each representing 20 discrete slices along the y-axis. Our method computes comparatively better segmentation in the challenging region below the large pial-vessel where image contrast is low due to occlusion.

### Graph-based modeling of cerebral vasculature

Vascular segmentation enables many important applications. Here, we are interested in graph-based modeling for brain vasculature [5, 28–30]. In this work, we propose a pipeline for graph extraction [Fig.1(A)], where we first compute the 3D segmentation using the method presented in this paper, followed by graph extraction using the framework recently proposed by Damseh et. al [18]. For each of the segmentation maps in Fig.3, including the ground truth segmentation, we compute the vascular graphs and present the comparative result in Fig.6. The qualitative results in Fig.6(A) demonstrate the graph computed from our segmentation to be more similar to the graph from the ground truth, especially for deeper vessels. This is in-line with our observation in Fig.3, where our method demonstrates comparatively better segmentation, particularly for deeper vessels. We also compute four basic metrics for quantitatively comparing the extracted graphs. First is the total number of vascular segments in the graph network. Second, the number of dangling segments, i.e. segments disconnected at on end or both ends. Generally there should not be any dangling segments except at the borders of the graph. Third, the number of short vascular segments, which we consider to be less than 6*μ*m. Finally, the number of incorrect bifurcations in the graph network. At bifurcation junctions, a vessel divides into two sub-vessels, and generally does not divide into more than two sub-vessels. If the extracted graph contains junctions where a vessel divides into three or more sub-vessels, we consider it to be an incorrect bifurcation. We compare these metrics among the extracted graphs in Fig.6(B), and find that the performance of our method resembles most closely to the ground truth. We also compute the graphs using the segmentation maps from Fig.5, and present the qualitative comparison in Fig.S2, empirically demonstrating satisfactory graph extraction performance up to a depth of 600*μ*m. We thus demonstrate that our segmentation method is suitable for large-scale vascular modeling and subsequent graph extraction.

**Figure 6.**
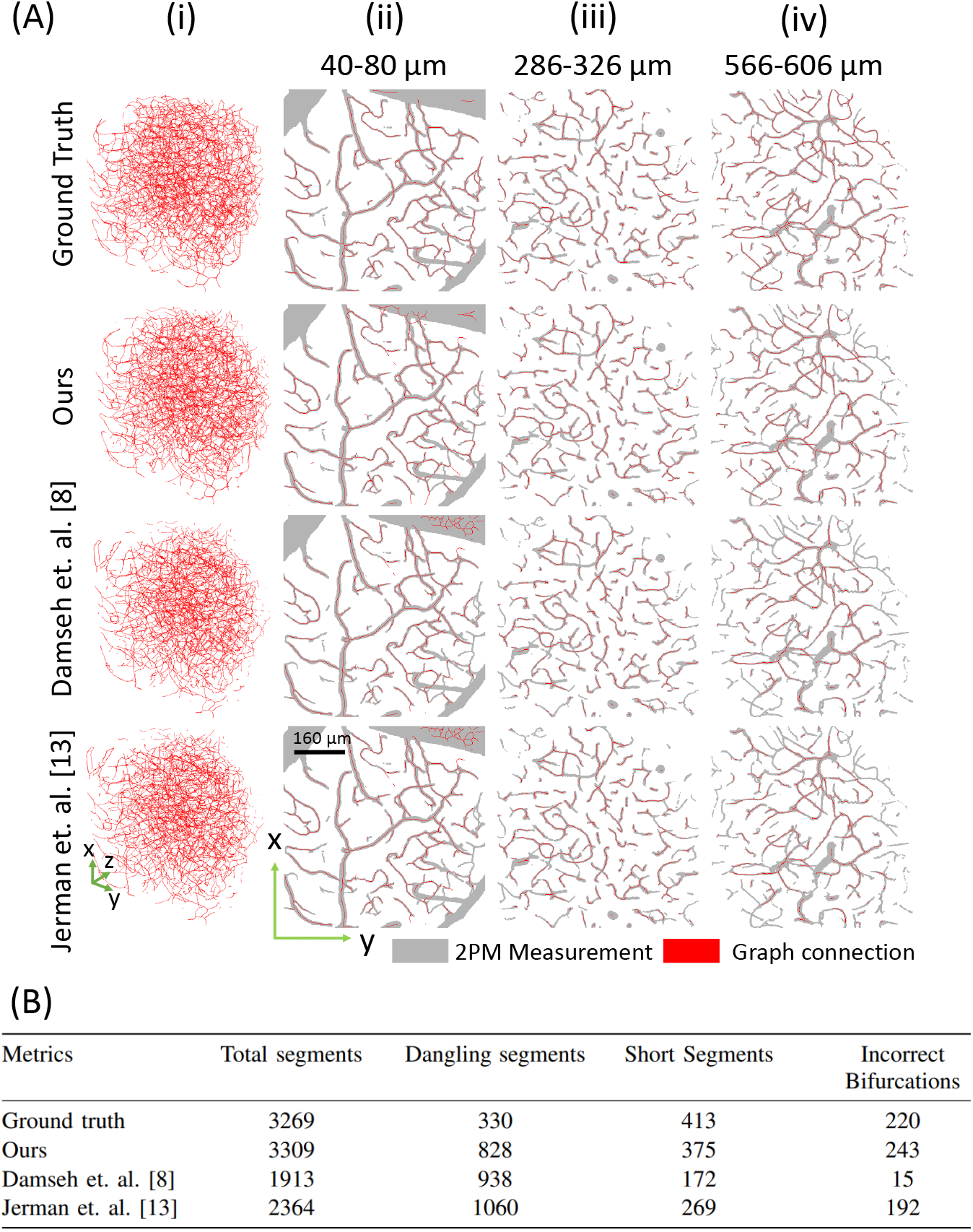
Graph extraction from 3D segmentation. The mathematical graph of the vasculature, comprising of nodes connected via edges, was computed from the segmentations in Fig.3. (A) Qualitative comparison of graph extraction performance. (i) 3D view of the graphs, depicted as vascular center lines in the volume. (ii-iv) MIPs of graphs overlaid on 2PM measurement, each MIP representing 20 discreet slices along z-axis. Graph extraction from our segmentation is qualitatively better compared to other methods, especially at increased depth. (B) A comparison of metrics demonstrates that the graph computed from our segmentation is quantitatively most similar to the graph from the ground truth segmentation.

## 3 Discussion

We propose and experimentally demonstrate a novel method for segmentation of 2PM angiograms, with the goal of large-scale cerebrovascular modeling. This new strategy enables processing of much larger angiograms compared to existing methods with significantly faster computation speed, by leveraging recent advances in deep learning. In addition, our deep neural network is able to segment angiograms from multiple 2PM imaging systems without retraining, and this flexibility shows its potential to be used as a general 3D segmentation tool for large-scale angiograms obtained using any 2PM imaging setup. In light of our goal of graph-based modeling of cerebral vasculature, we compute vascular graphs from binary segmentation, using a technique recently developed by one of our co-authors [18]. We observe that improved segmentation using our method led to better vascular graphs for large 2PM angiograms. This has important implications since existing graph extraction pipelines do not demonstrate adequate accuracy, and have to be followed up by significant manual correction as a post-processing step [30]. This human annotation can quickly become infeasible as the angiograms scale to greater sizes and quantities. It is therefore desirable to have a method for accurate graph computation which can minimize, if not completely eliminate, the use of manual correction. Towards this end, we have presented a modular approach for graph computation, where the challenging 2PM vascular segmentation has been decoupled from graph extraction. This gives us the ability to optimize each of these two steps independently.

While our method was able to demonstrate significantly deeper segmentation compared to existing techniques, it still has several limitations. Our method was unable to accurately segment vasculature beyond 600*μ*m within the brain tissue. This is partly due to the limitation of 2PM to capture angiograms with sufficient SBRs much beyond this depth, and also due to the unavailability of accurate ground truth for deeper angiograms. Effective segmentation for deeper vasculature might be achieved by employing ground-truth data with greater depth, coupled with more intelligent semi-supervised learning, involving e.g. active contours [31], in addition to the TV regularization used in our work. Another limitation is that angiograms from different setups have to undergo manual histogram equalization before being segmented by our network. This involves linear scaling to make the voxel distribution of new angiograms similar to those on which the network has been trained. Further work may look to automate the process. In general, more advanced domain adaptation techniques [32, 33] may be incorporated to further improve the generalizability. Although we demonstrate improved segmentation performance in the low-contrast region under large pial vessels compared to existing methods, the segmentation still suffers from artifacts and obvious false negatives. Further work may look to improve the performance in such regions, either on the acquisition end by employing better fluorophores, or by using a pre-processing method to enhance the contrast of the vasculature under pial vessels, prior to segmentation. Despite these limitations, we demonstrate state-of-the-art performance for vascular segmentation of large-scale 2PM angiograms. We believe that this work paves the way towards large-scale cerebrovascular modeling and analysis.

## 4 Materials and method

### 4.1 Data preparation

2PM angiograms were acquired on two different imaging systems for various mice specimen (*n* = 5 for system 1, and n = 1 for system 2). For training and quantitative evaluation, we used data only from the first imaging setup, while data from the second setup was used for qualitative demonstration of the generalizability of our approach.

The dataset from imaging system 1 has been previously published by Gagnon et. al. [5, 30], and it’s preparation is detailed as follows. All experimental procedures were approved by the Massachusetts General Hospital Subcommittee on Research Animal Care. C57BL/6 mice (male, 25–30 g, *n* = 5) were anesthetized by isoflurane (1–2% in a mixture of *O*_2_ and air) under constant temperature (37° *C*). A cranial window with the dura removed was sealed with a 150-m-thick microscope cover-slip. During the experiments, a catheter was used in the femoral artery to monitor the systemic blood pressure and blood gases and to administer the two-photon dyes. During the measurement period, mice breathed a mixture of *O*_2_, and air under the 0.7–1.2% isoflurane anesthesia. Structural imaging of the cortical vasculature was performed using a custom built two-photon microscope [34] after labeling the blood plasma with dextran-conjugated fluorescein (FITC) at 500 nM concentration. Image stacks of the vasculature were acquired with 1.2 × 1.2 × 2.0 μm voxel sizes under a 20× Olympus objective (NA= 0.95). Data was digitized with a 16 bit depth. A total of five angiograms were acquired on this setup, each from a distinct specimen Fig.S3(A-E), and were divided into training and testing angiograms in a ratio of 80-20% respectively, i.e. four angiograms were used for training [Fig.S3(B-E)], while one was used for testing and evaluation [Fig.S3(B-E)]. The ground-truth segmentation was prepared by human annotators using custom software.

For imaging system 2, the dataset has a similar preparation process for the live specimen, however, it was acquired on a different imaging system and different mouse whose details are as follows. All experimental procedures were approved by the BU IACUC. We anesthetized a C57BL/6J mouse by isoflurane (1–2% in a mixture of O_2_ and air) under constant temperature (37°*C*). A cranial window with the dura intact was sealed with a 150-m-thick microscope cover-slip. During the measurement period, mice breathed a mixture of O_2_, and air under the 0.7–1.2% isoflurane anesthesia. The blood plasma was labeled using dextranconjugated fluorescein (FITC) at 500 nM concentration. Imaging was performed using a Bruker two-photon microscope using a 16× objective (NA=0.8) with voxel size 1.58 × 1.58 × 2.0 μm. Data was digitized with 12 bit depth. One angiogram was acquired on this setup Fig. S3(F), and was used as test data the generalization capability of our network.

### 4.2 Data preprocessing

Adequate pre-processing on test-data was found to be critical for good network generalization. Here, we present a two-step preprocessing method that consists of histogram-scaling, followed by noise removal, for improved segmentation.

Since our two imaging setups have different detector bit-depths, their respective angiograms also differed with respect to the scale of voxel-intensities Fig.S4(A,C). Since our DNN learns a maximum likelihood function for mapping the input angiograms to the desired 3D segmentation maps, given the training data, it is important that the test angiogram from any imaging system is on a similar intensity scale as the training data. Noticing that the histograms from both setups are similar in shape [Fig.S4(A,C)], but different with respect to intensity scale; we perform linear scaling on data from setup 2 by multiplying it with a non-negative scaling factor. The scaling factor is chosen such that after scaling, the intensity histogram of the angiogram from setup 2 becomes similar in scale to the intensity histogram of the angiogram from setup 1. This procedure has been depicted in Fig.S4(B,D). The scaling factor for our case, was empirically chosen to be 16. In the case of applying this approach to an angiogram from a different imaging setup, the procedure will be very similar. This new angiogram would have to be multiplied with a non-negative scaling factor, which is empirically chosen such that the voxel-intensity histogram of the scaled angiogram becomes similar in scale to that of an angiogram from setup 1.

A well-known challenge inherent to 2PM is the degradation of signal with imaging depth [Fig.2, Fig.S5(A)]. Segmentation on such an angiogram using our trained DNN has significant artifacts, even after linear scaling. Here we propose a simple yet effective method to reduce this depth-dependent noise. We subtracted from each 2D image in a 2PM 3D stack, it’s median value. This visibly improved the signal quality by suppressing the background noise, especially in deeper layers [Fig.S5(B)]. The angiogram was further improved by applying 3D median filter with a kernel of size 3 voxels [Fig.S5(C)]. This preprocessing method improved the segmentation of the deep vasculature, and made individual vessels more distinguishable [Fig.S6]. However, this method was observed to decrease segmentation quality in the shadowed region under large pial vessels where the measurement contrast is comparatively weak. A locally adaptive pre-processing method that could overcome this limitation may be a potential direction of future work.

### 4.3 Deep neural network design and implementation

Our DNN architecture is based on the well-known V-net [35], however we significantly modified the original framework for large-scale 2PM vascular segmentation [Fig.S7]. The network is end-to-end 3D for fast computation, as opposed to 2D slice-wise techniques, and consequently also takes into account the 3D context for improved segmentation. It has an encoder-decoder framework for learning vascular features at various size-scales, and high resolution feature forwarding to retain high-frequency information. We incorporate batch-normalization after each convolution-layer to improve generalization performance and convergence speed. Our network processes 3D input patches with outputs of the same size. Patch based processing enables the segmentation of arbitrarily large volumes. Our large patch-size compared to existing methods, coupled with a lightweight network, help to significantly accelerate computation speed. For the training process, the training data is divided into patches of 128 × 128 × 128 voxels, with an overlap of 64 voxels along all axes. Each training iteration processes a batch of 4 patches chosen randomly from the training data. We use Adam optimizer to train our network with a learning rate of 10^-4^ for about 100 epochs, which takes approximately 4 hours on a TitanXp GPU. For testing, the angiogram is divided into patches of 128 × 128 × 128 voxels and segmentation is performed on each patch separately, after which they are stitched together to get the final segmented angiogram. The division of the acquired data into training and testing datasets has been described in Section 4.1.

### 4.4 Loss function design

During the training process of a DNN, a loss function is optimized via gradient descent or any of its variants. The loss itself is a function of the network output, and is chosen by the user to impart desired characteristics to the DNN output by guiding the training process. In this problem, we initially experimented with binary cross entropy (BCE) loss, as it is known to promote sparsity in the output [36], which is desirable for vascular segmentation. However, severe class-imbalance in our data rendered BCE ineffective as a loss function, and the DNN converged to a nearly zero-solution, i.e. almost all voxels were classified as background. Class-imbalance is the situation when one class significantly outnumbers the others in the training data, causing a preferential treatment by the learning algorithm towards the abundant class. In our case, the negative class consisting of background-voxels was significantly more abundant than the positive class, and constituted 96% of the total voxels in the training data. This resulted in the significant number false negatives in the DNN predictions using BCE. In order to overcome this challenge, we incorporated a variant of BCE loss with class balancing [25, 37], 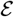, defined as

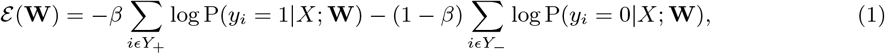

where **P** is the probability of obtaining the label *y_i_* for the *i*^th^ voxel, given data *X* and network weights **W**. *β* and (1 – *β*) are the class weighting multipliers, defined as 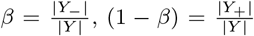, where *Y*_+_ is the set of positive (vessel) labels, and *Y*_-_ is the set of negative (background tissue) labels, *Y* being the set of all voxels, both vessel and background. In this loss, we essentially weigh down the negative class, and give a greater weight to the positive class, and the assigned weight depends on the fractions of vessel and background voxels in the volume, respectively. Note that *β* is not a tunable hyperparameter here, rather, it’s value is determined by the training data in every iteration. This balanced BCE loss significantly improved training in the presence of severe class imbalance.

Merely using the balanced BCE loss described above was found to be insufficient to provide satisfactory generalization performance. One way to improve generalizability is to use training data from various different imaging setups. However, manually annotating many large-scale angiograms for this purpose would have been prohibitive due to the associated time and cost. We therefore took a different approach and employed TV regularization in the loss function, which improved generalization in the presence of limited and noisy training data [38]. For this purpose, we added a regularization term to the loss function, which penalizes the total variation of the network output [21]. Our TV-loss, 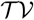, is defined as

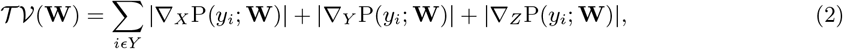

where ∇_*X*_, ∇_*Y*_, and ∇_*Z*_ are 3D Sobel operators for computing TV [39]. 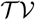 when added to the balanced BCE loss, decreases the model dependence on the ground truth data, helping generalization. TV is known to promote sparsity and piece-wise continuity in solutions, which are suitable priors for vascular segmentation. The addition of TV imparted de-noising property to the network, such that no post-processing was required on the outputs after segmentation; and improved the generalization performance [Fig.S8].

Finally, we also regularize our loss function by adding a penalty on the *l*_2_-norm of the network weights **W**. This is called weight decay, and is known to encourage the network to learn smooth mappings from the input angiogram to the output segmentations, reducing over-fitting and improving generalization. The final form of our loss function 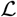 is thus

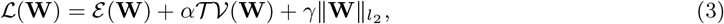

where *α* and *γ* are tunable parameters, whose values were empirically found to be 5 × 10^-9^ and 0.01 respectively for best performance. We present how different levels of TV regularization impact the segmentation performance in Fig.S9, and demonstrate the range for optimal value of *α*. Similarly, we also show segmentation performance as a function of *γ*, and present the optimal range for weight decay in Fig.S10.

### 4.5 Segmentation evaluation metrics

Accuracy = *TP* + *TN*/(*TP* + *TN* + *FP* + *FN*), Jaccard index = *TP*/(*TP* + *FP* + *FN*), Dice coefficient = 2*TP*/(2*TP* + *FP* + *FN*), Specificity = *TN*/(*FP* + *TN*), and Sensitivity = *TP*/(*TP* + *FN*). Here, TP (True Positive) is the number of correctly classified vessel voxels, TN (True Negative) is the number of correctly classified background voxels, FP (False Positive) is the number of background voxels incorrectly labeled as vessels, and FN (False Negative) is the number of vessel voxels incorrectly labeled as background.

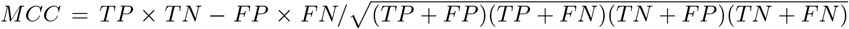, which measures the linear correlation between the ground truth and predicted labels and is a special case of the Pearson correlation coefficient.

HD among two finite point sets can be defined as HD(*A, B*) = max(*h*(*A, B*), *h*(*B, A*)), where 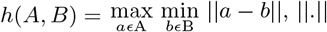 being any norm e.g. Euclidean norm.

LC is defined as LC(*S, S_G_*) = #((*g*(*S*) ⋂ *S_G_*) ∪ (*S* ⋂*g*(*S_G_*)))/#(*g*(*S*) ∪ *g*(*S_G_*)), where *S* and *S_G_* are the predicted and ground truth segmentation, respectively, *g*(.) is an operator that computes the 3D vascular skeleton from an input segmentation in the form of a graph of nodes and edges, using the method in [18], and #(.) measures the cardinality of an input set in terms of number of voxels.

## 5 Acknowledgements

**General:** We thank David Kleinfeld (UCSD), Timothy W. Secomb (University of Arizona), Chris B. Schaffer (Cornell University), and Nozomi Nishimura (Cornell University) for helpful discussions, Yujia Xue (Boston University) and Yunzhe Li (Boston University) for the help in running experimental code, and Alex Matlock (Boston University) for reviewing the manuscript.

## Author contributions

W.T. developed and implemented software for segmentation and wrote the paper. S.K. helped with data preparation, deep learning design, paper writing and performed results analysis. J.Z. contributed to deep neural network design and implementation. F.T. performed ground truth annotation. X.C. contributed to experimental design, results analysis and paper writing. B.S.L. performed data acquisition on setup 2. A.S. implemented the graph extraction algorithm on the BU cluster. R.D. developed graph extraction software and helped with it’s implementation. F.L. supervised graph extraction from vascular segmentation. S.S. helped with experimental design and analysis of data from setup 1. D.B. and L.T. initiated, supervised, and coordinated the study, and help write the manuscript.

## Data availability

All data needed to reproduce and evaluate the results in this paper has been made publicly available at https://github.com/bu-cisl/2PM_Vascular_Segmentation_DNN.

## 6 Supplementary material

**Figure S1.**
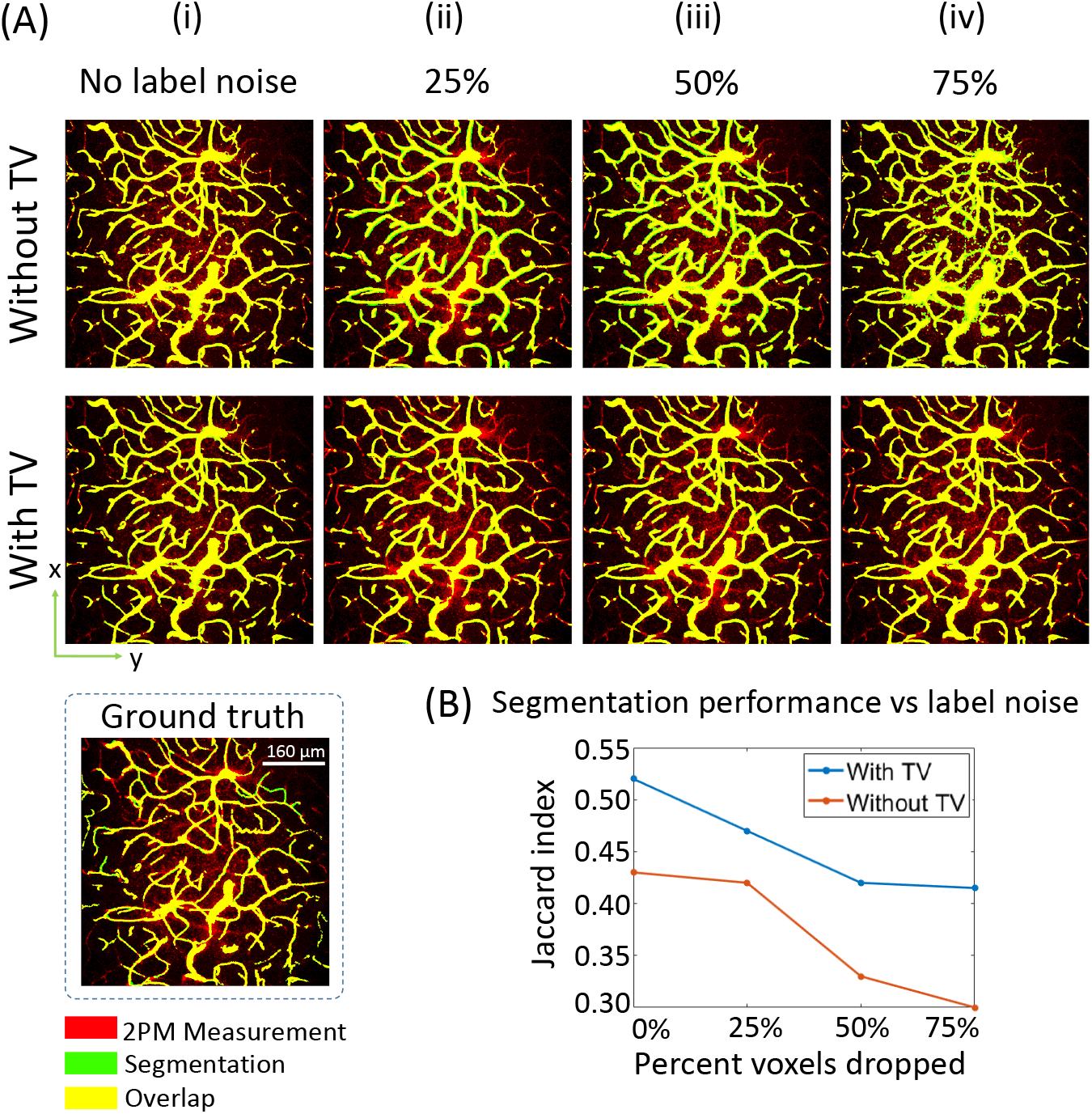
Robustness of the proposed learning scheme to label noise. We train our DNN with various levels of label noise, and demonstrate that adding TV-regularization improves robustness of the segmentation method to label noise. To add label noise, the foreground labels (vessels) were first dilated with a sphere of radius 2. Then a certain percentage of the foreground voxels was selected randomly with uniform probability, and were dropped to zero, such that they now represented background. This method of adding noise was especially chosen to make the edges of vessels ambiguous in the ground truth, as vascular edges are most prone to mislabeling.(A) Qualitative comparison of segmentation performance with various levels of label noise (MIPs 566 — 606μm). The top row represents results from our DNN without TV-regularization (α = 0), while the bottom row represents our DNN with TV-regularization. (i) The baseline performance where no label noise has been added to the ground truth. (ii-iv) Label noise is progressively increased from 25% to 75%. The results without TV-regularization deteriorate significantly, while the DNN with TV-regularization is visibly more robust, even in the case of 75% label noise. (B) The quantitative analysis also supports the qualitative comparison between TV and no-TV segmentation; and demonstrates that TV-regularization imparts robustness to label noise.

**Figure S2.**
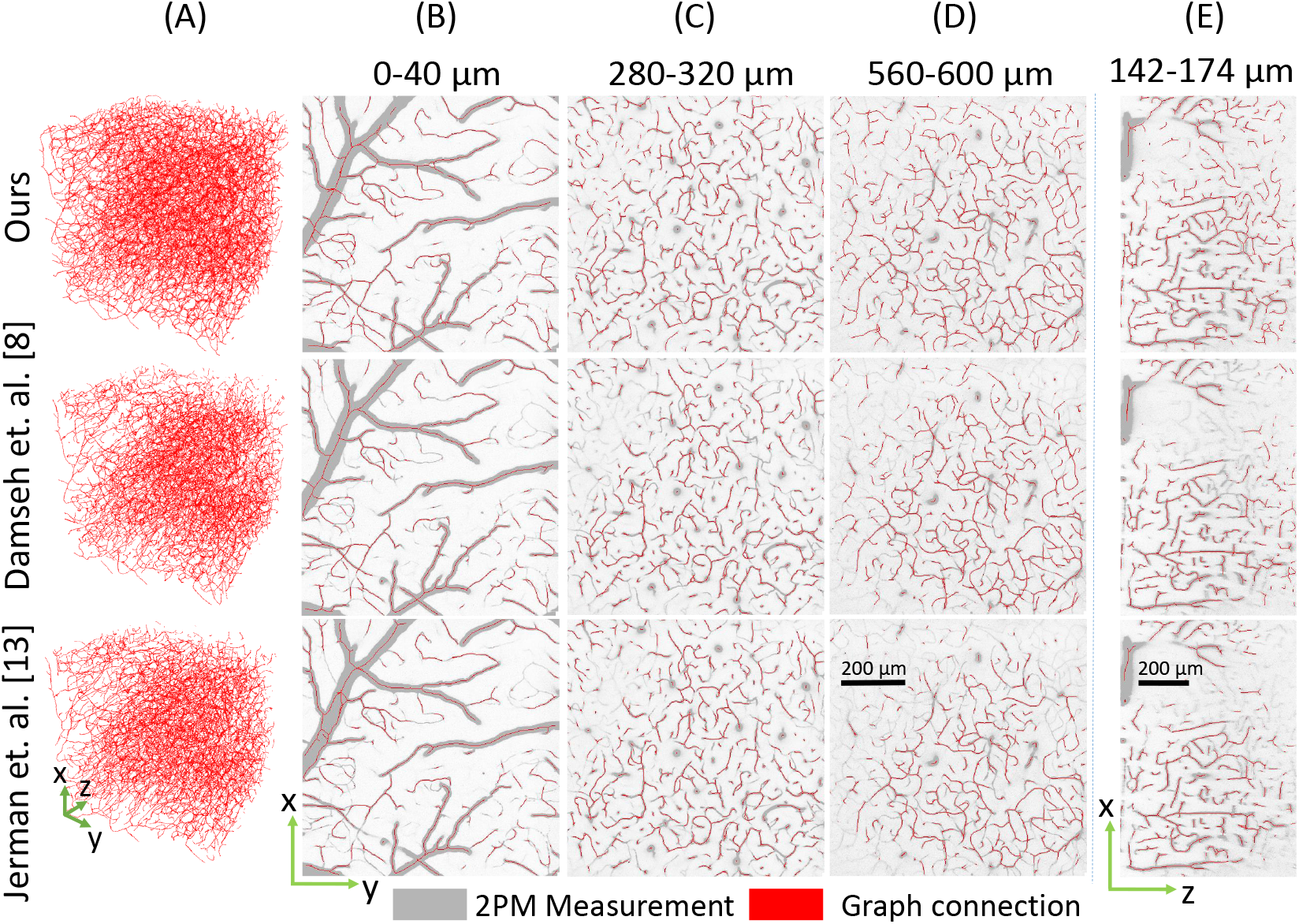
Graph extraction from the 3D segmentation map. The mathematical graph of the vasculature was computed from the segmentation, comprising of nodes connected via edges. (A) 3D view of the graphs, depicted as vascular center lines in the volume. (B-D) MIPs of graphs overlaid on 2PM measurement, each MIP representing 20 discreet slices along z-axis. (E) Longitudinal x-z MIP overlays, each MIP representing 20 discreet slices along y-axis. Graph extraction from our segmentation is qualitatively better compared to other methods, especially below the large pial vessel where measurement contrast is low, and for deep vasculature.

**Figure S3.**
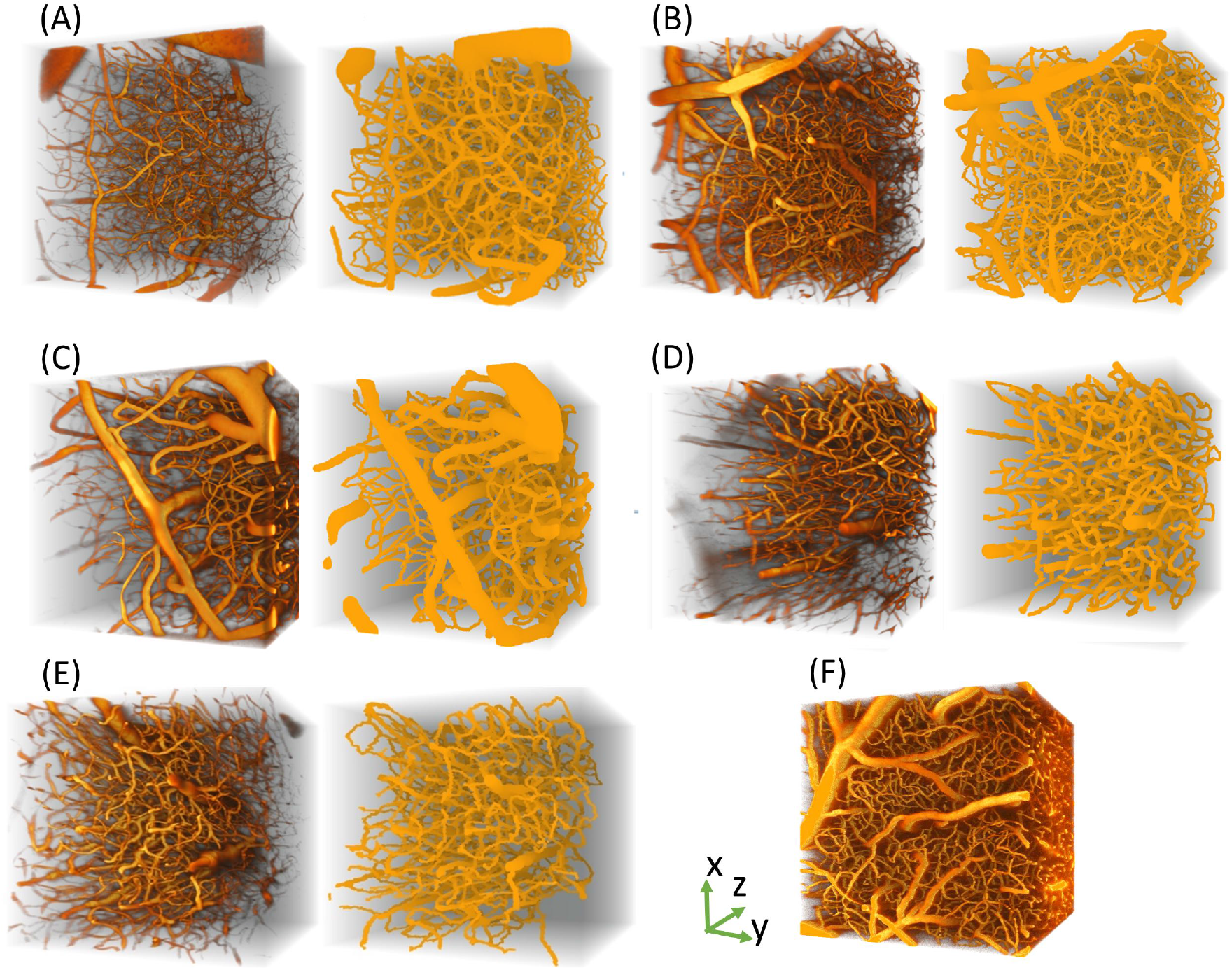
Training and testing data used in our experimentation. (A-E) 2PM measurements and annotated ground truth segmentation pairs for 5 angiograms from setup 1. (F) 2PM measurement from setup 2.

**Figure S4.**
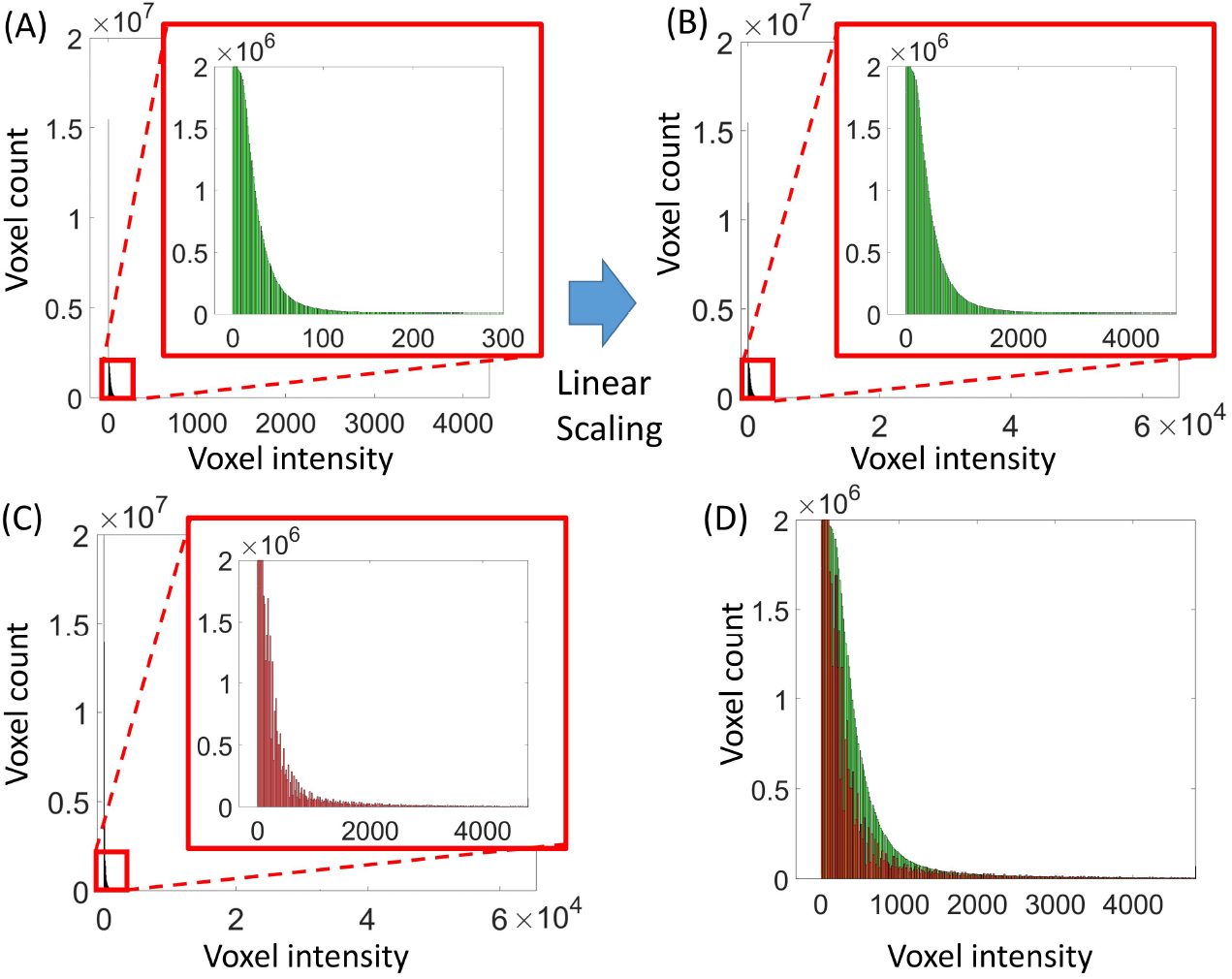
Data pre-processing: intensity scaling. (A) Histogram for the angiogram from setup 2. (B) Histogram in (A) after linear scaling. Scaling is performed on data by multiplying the angiogram with a constant factor to make the intensity scale similar to data from setup 1. (C) Histogram for the test mouse from setup 1. The intensity scale is significantly different from (A) due to difference in bit-depth of camera between setups 1 and 2. (D) Overlay of (B) and (C). The intensity scales between data from setup 1 and 2 become similar after linear scaling on data from setup 2.

**Figure S5.**
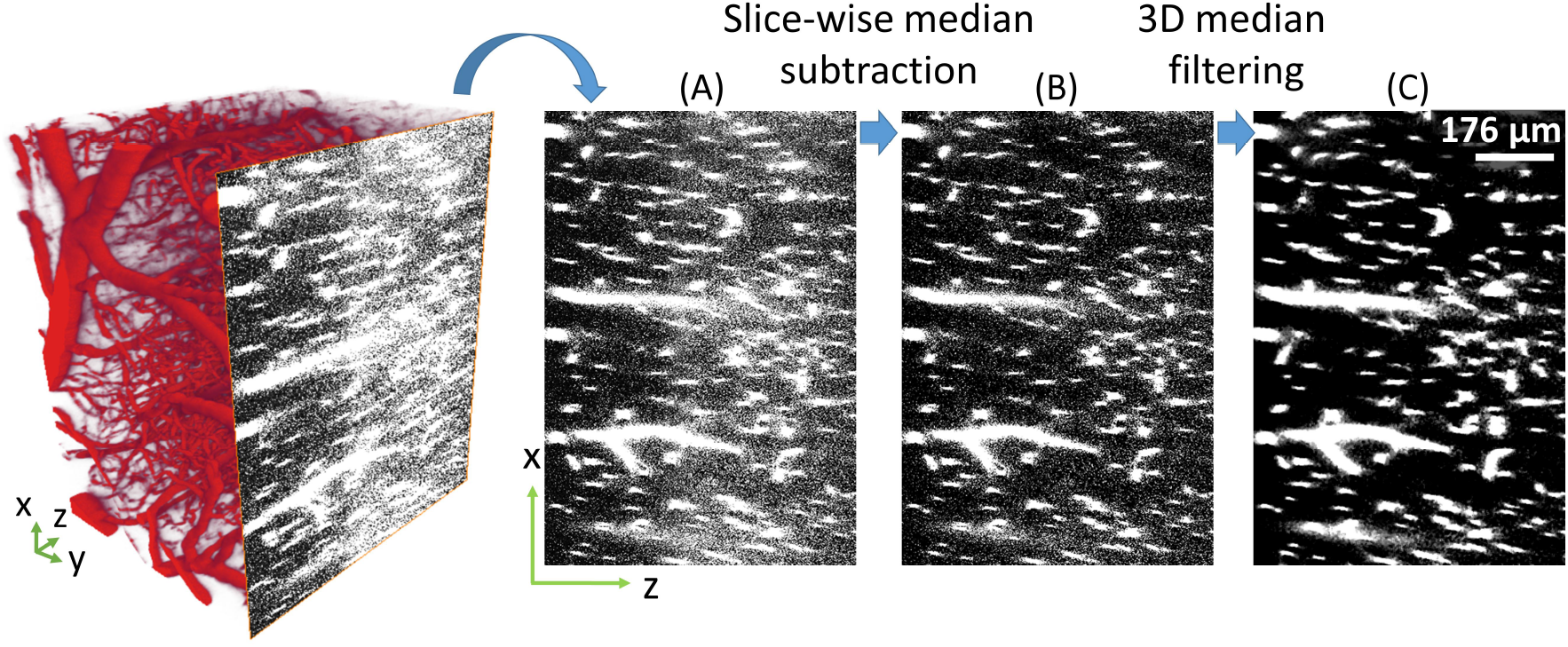
Data pre-processing: denoising. (A) 2PM inherently suffers from signal degradation with imaging depth. (B) Subtracting from each 2D image in the 3D stack, it’s median value, visibly improved the quality of the angiogram. (C) 3D median filtering on the angiogram with a 3 × 3 × 3 window significantly reduced background noise.

**Figure S6.**
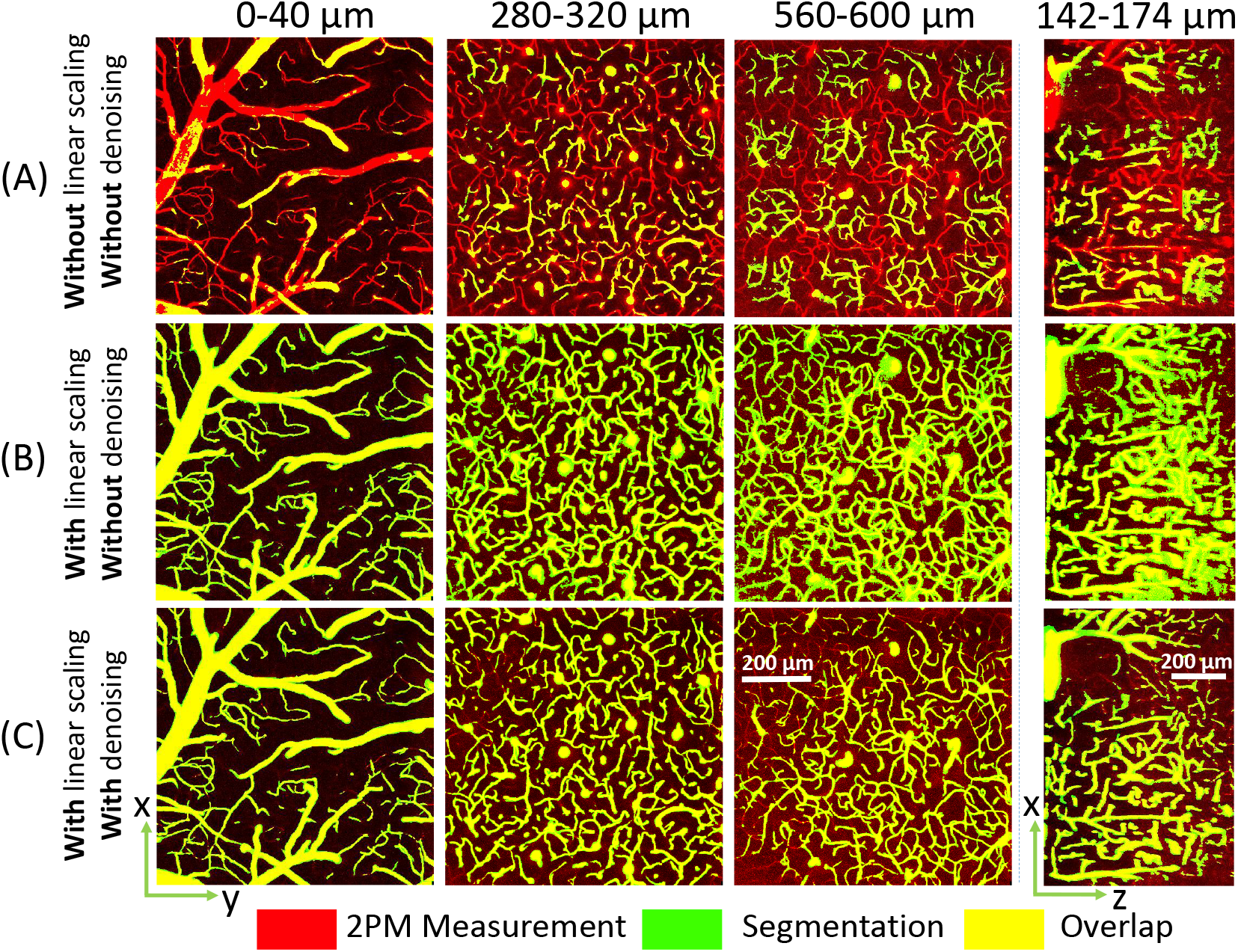
Effect of different steps of preprocessing on segmentation quality. (A) Without any preprocessing on data from setup 2, the segmentation suffers from significant artifacts. (B) Applying linear scaling improves the number of vessels recovered in the segmentation. However, there is a significant number of false positives on vessel boundaries, leading to many adjacent vessels being joined together in the segmentation map. (C) Application of slice-wise median subtraction and 3D median filtering, jointly referred to as ‘denoising’, further improves the segmentation. Even though the challenging region behind the large pial vessel does contain missed vessels in this case, we do achieve significantly better distinction of vessels in the segmentaiton. Note that in all cases above, we use the DNN trained with the loss described in Eq.(3).

**Figure S7.**
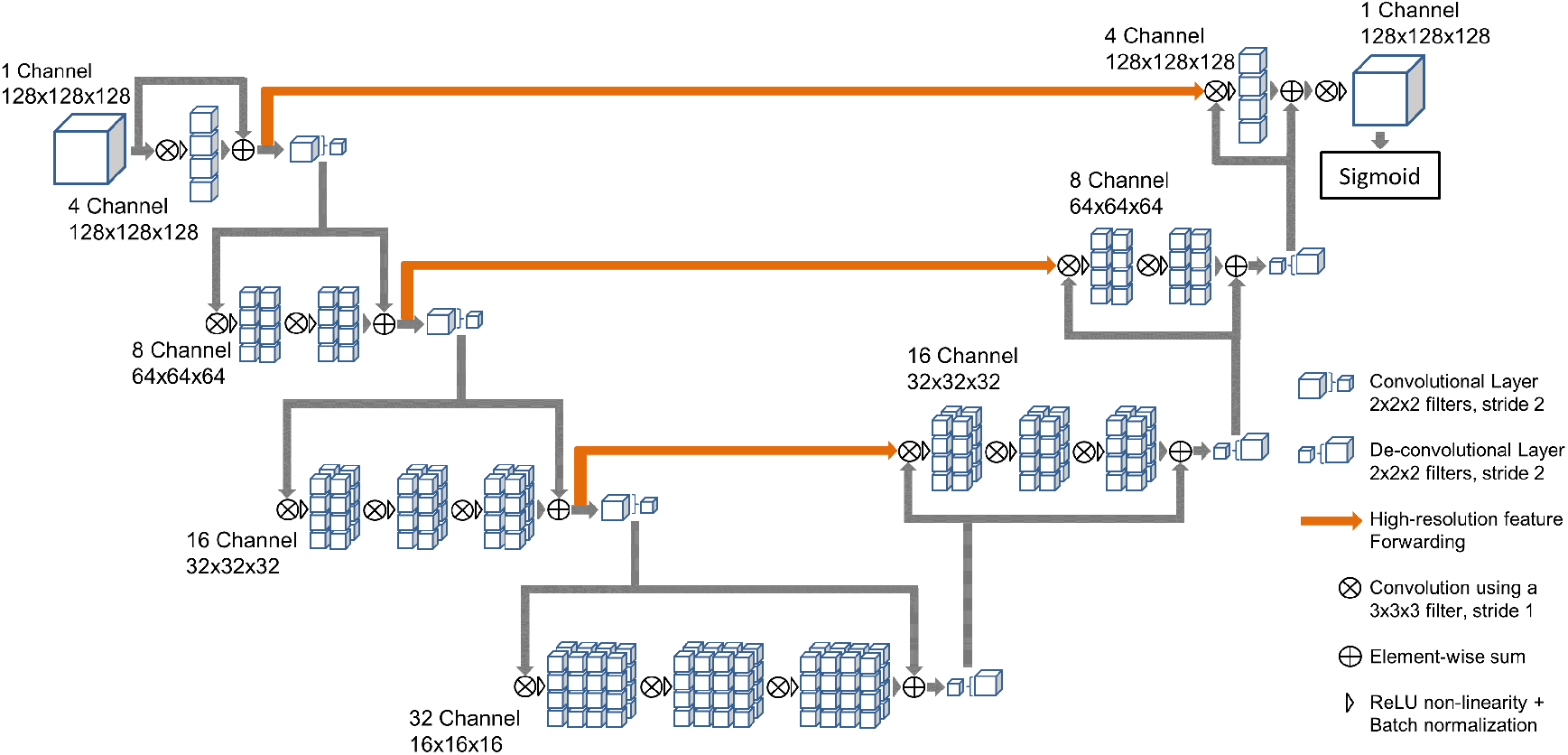
Deep neural network architecture. Our network is designed to be fast and generalizable with good segmentation accuracy. The network is end-to-end 3D taking into account 3D context. Skipped connections forward high-resolution features. Batch normalization improves generalizability and convergence speed, and the number of layers and weights are chosen so as to minimize processing time and over fitting, while maintaining segmentation accuracy.

**Figure S8.**
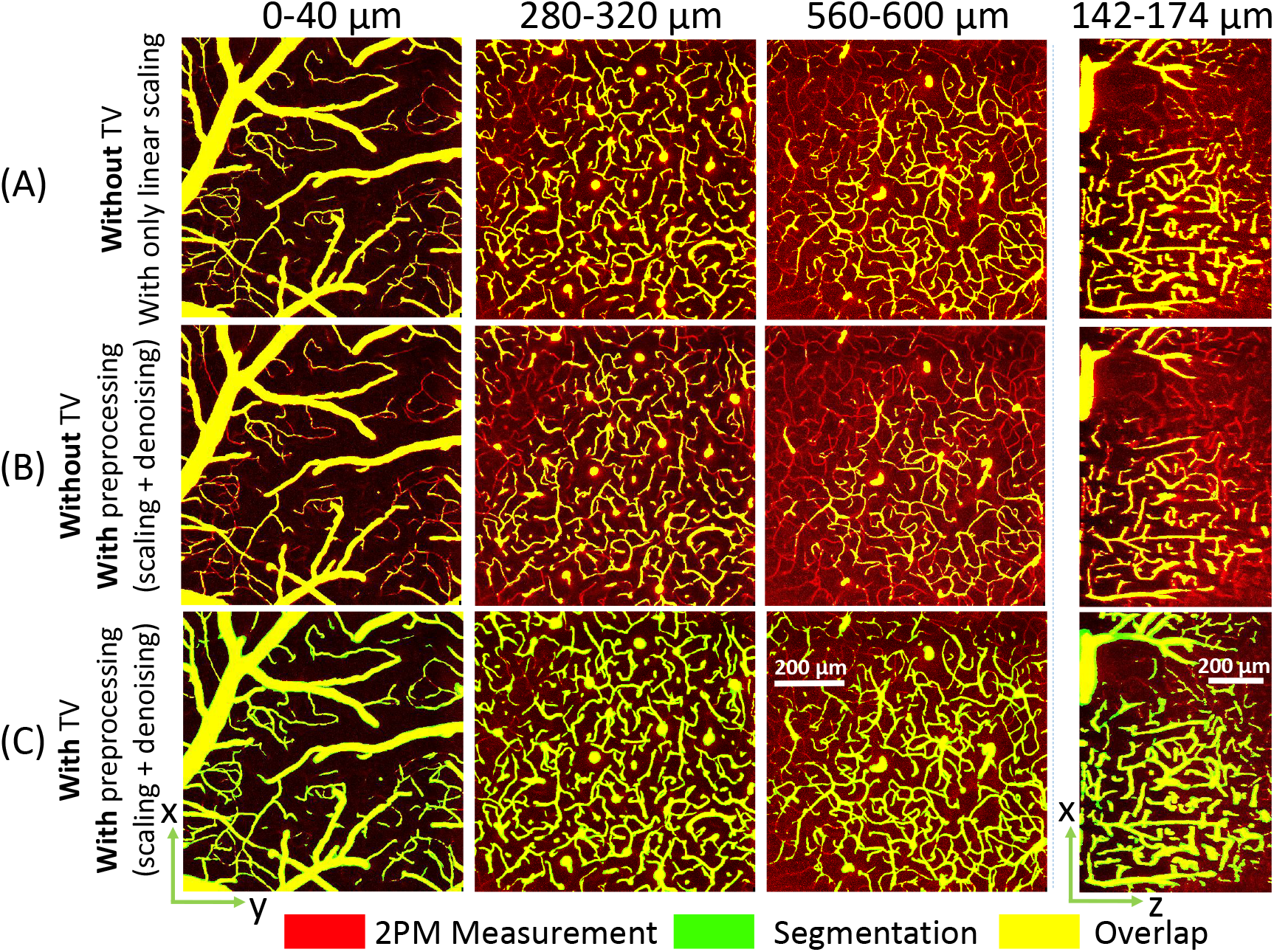
Ablation of TV and preprocessing for segmentation. Here we compare the performance of our DNN with and without TV regularization in the loss function, and two levels of preprocessing. (A) Here only linear scaling is performed on the input angiogram before segmentation with a DNN trained without TV. There are a significant number of missed vessels especially in the region below the large pial vessel. (B) Here complete preprocessing is performed on the input angiogram including linear scaling and denoising, before segmentation with a DNN trained without TV. (C) Here TV is added to the DNN loss, in addition to performing preprocessing on the input angiogram, and we see that the segmentation quality is significantly improved. Since this segmentation is on an anigogram from setup 2, it also points towards improved generalization as a result of TV regularization. It is noteworthy that without any preprocessing at all, i.e. no linear scaling and no denoising, the network output without TV regularization was majorly all zero.

**Figure S9.**
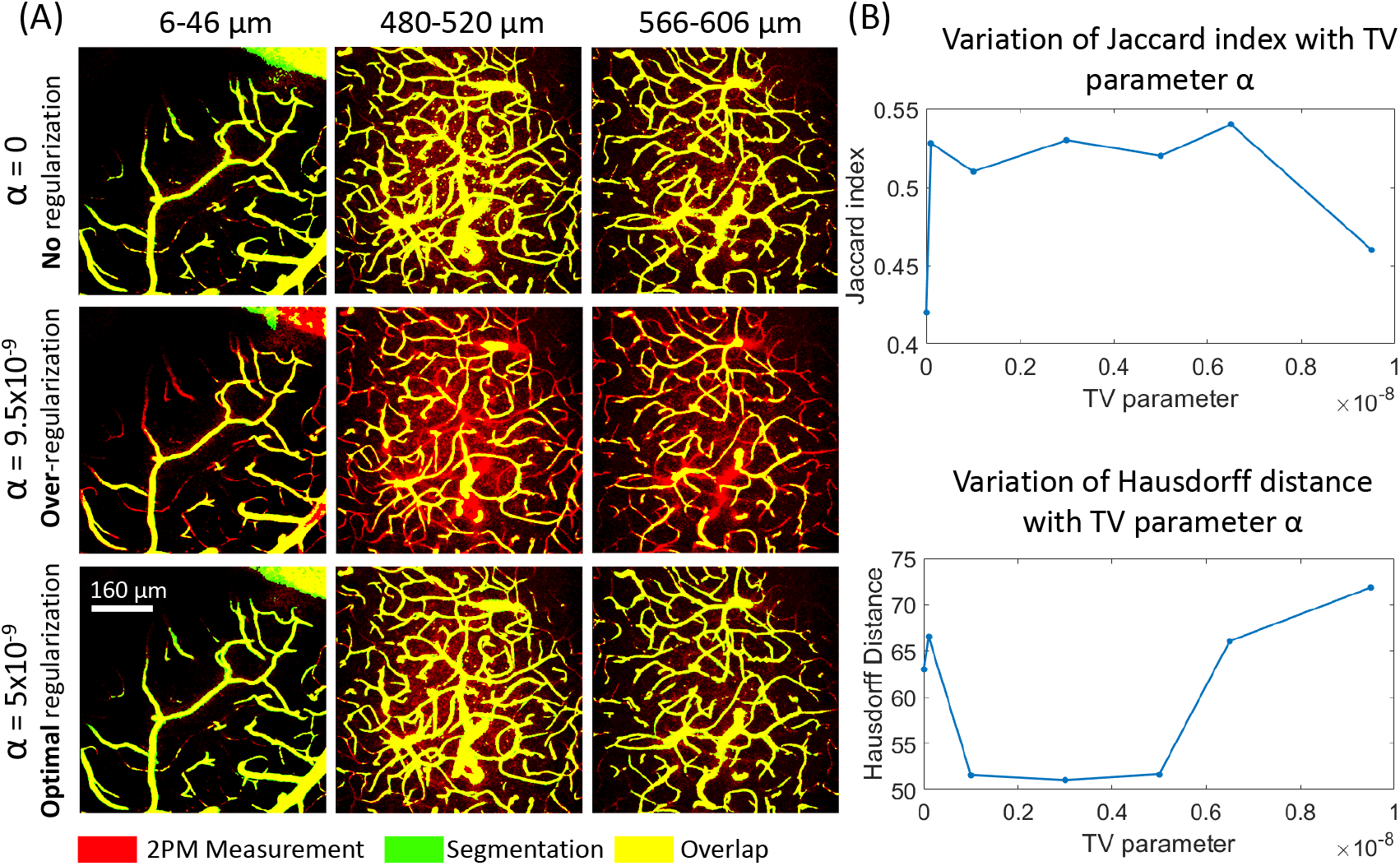
Effect of total variation (TV) regularization on segmentaion performance. (A) Without any regularization (α = 0), we observe visible noise in the background, and nonsmooth vascular boundaries. When the regularzation parameter is set very high, the segmentation has many missed vessels as the DNN tries to minimize TV. In the optimal TV range, the segmentation quality is the best, both quantitatively and qualitatively. (B) We present the quantitative segmentation quality as a function of the TV parameter α, using Jaccard index, and Hausfdorff distance, as metrics. We find the optimal range of α to be about 1 × 10^-9^ to 5 × 10^-9^.

**Figure S10.**
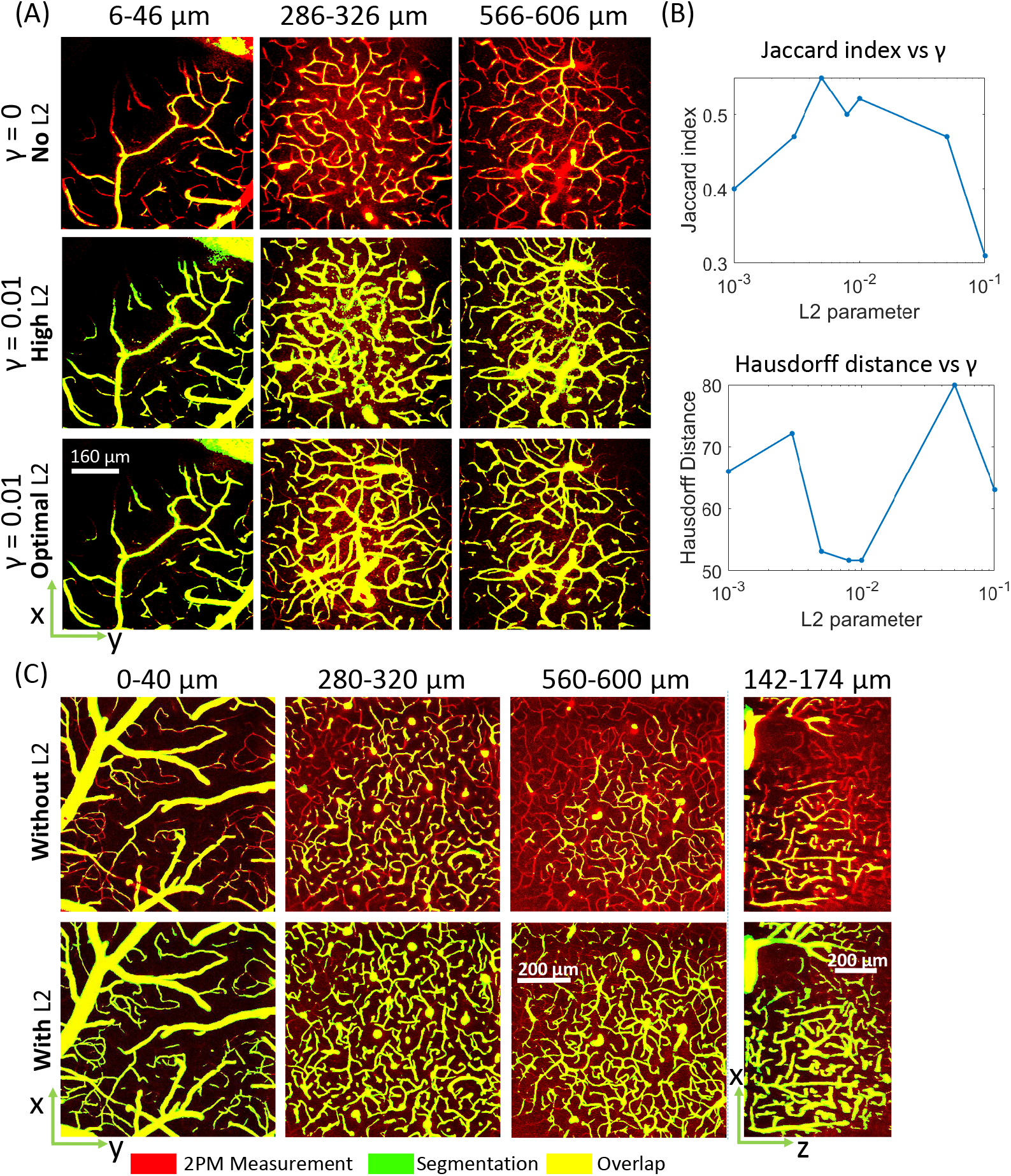
Effect of L2 regularization (weight decay) on segmentaion performance. (A) Without weight decay (*γ* = 0), we observe sub-par segmentation performance and a significant number of missed vessels. When *γ* is set very high, the weights are unable to optimize, resulting in bad segmentation performance. In the optimal range of *γ*, we obtain best segmentation performance. (B) We present the quantitative segmentation quality as a function of weight decay, using Jaccard index, and Hausfdorff distance, as metrics. We find the optimal range of *γ* to be about 5 × 10^-3^ to 1 × 10^-2^. (C) Here we compare segmentation maps for the angiogram from setup 2, with and without weight decay. We see that weight decay significantly improves the segmetnation quality, and thus improves the generalization ability of the DNN.

